# Selective loss of CD107a TIGIT+ memory HIV-1-specific CD8+ T cells in PLWH over a decade of ART

**DOI:** 10.1101/2022.07.13.499924

**Authors:** Oscar Blanch-Lombarte, Dan Ouchi, Esther Jiménez-Moyano, Julieta Carabelli, Miguel Angel Marin, Ruth Peña, Adam Pelletier, Aarthi Talla, Ashish Sharma, Judith Dalmau, José Ramón Santos, Rafik-Pierre Sékaly, Bonaventura Clotet, Julia G. Prado

## Abstract

The co-expression of inhibitory receptors (IRs) is a hallmark of CD8+ T-cell exhaustion (Tex) in people living with HIV-1 (PLWH). Understanding alterations of IRs expression in PLWH on long-term antiretroviral treatment (ART) remains elusive but is critical to overcoming CD8+ Tex and designing novel HIV-1 cure immunotherapies. To address this, we combine high-dimensional supervised and unsupervised analysis of IRs concomitant with functional markers across the CD8+ T-cell landscape on 24 PLWH over a decade on ART. We define irreversible alterations of IRs co-expression patterns in CD8+ T cells not mitigated by ART and identify negative associations between the frequency of TIGIT+ and TIGIT+TIM-3+ and CD4+ T-cell levels. Moreover, changes in total, SEB-activated, and HIV-1-specific CD8+ T-cells delineate a complex reshaping of memory and effector-like cellular clusters on ART. Indeed, we identify a selective reduction of HIV-1 specific-CD8+ T-cell memory-like clusters sharing TIGIT expression and low CD107a that can be recovered by mAb TIGIT blockade independently of IFNγ and IL-2. Collectively, these data characterize with unprecedented detail the patterns of IRs expression and functions across the CD8+ T-cell landscape and indicate the potential of TIGIT as a target for Tex precision immunotherapies in PLWH at all ART stages.

## Introduction

The ART introduction has been the most successful strategy to control viral replication, transforming HIV-1 into a chronic condition. However, ART does not cure the infection, and treatment is required lifelong due to a stable viral reservoir, raising the need to find a cure for people living with HIV-1 (PLWH). A sterilizing or functional cure aims to eliminate or control HIV-1 in the absence of ART. In both scenarios, HIV-1-specific CD8+ T cells are likely to play an essential role as they have been widely recognized as a critical factor in the natural control of viral replication (1–5). Proliferative capacity, polyfunctionality, and ex vivo antiviral potency are features of HIV-1-specific CD8+ T cells associated with spontaneous viral control (6–9).

Although ART in PLWH normalizes the levels of CD8+ T cells and potentially preserves their functional characteristics (9–12), microbial translocation and continuous immune activation lead to long-term CD8+ T-cell dysfunction and exhaustion (Tex), a critical barrier for HIV-1 curative interventions (13–15). CD8+ Tex is defined by the persistent co-expression of inhibitory receptors (IRs) and the progressive loss of immune effector functions linked to transcriptional, epigenetic and metabolic changes (16–19). In HIV-1 infection, IRs are continuously expressed despite long-term suppressive ART (20–26) and have been associated with disease progression and immune status in PLWH (25,27–32). Thus, the expression of IRs is linked to diminished functionality and is a hallmark of CD8+ Tex in PLWH.

The interest in evaluating the blocking of IRs or immune checkpoint blockade (ICB) in PLWH as a therapeutic strategy to reverse CD8+ Tex increases (26,27). Over the last years, several studies supported the recovery of proliferative capacity, cell survival, and cytokine production of HIV-1-specific CD8+ T cells through ICB (33,34). The blockade of the PD-1/PDL-1 axis has been extensively studied, demonstrating the functional recovery of HIV-1-specific CD8+ T cells in PLWH (26,27). Moreover, alternative pathways to PD-1/ PDL-1, including LAG-3, TIGIT, and TIM-3, have been explored as candidates for ICB therapies for HIV-1 infection (33,35–37). Also, recent data support the combinatorial use of ICB to favour synergistic effects on the recovery of HIV-1-specific CD4+ and CD8+ T-cell function (38,39).

The unprecedented success of the clinical use of ICB in the cancer field (40,41) has prompted the clinical evaluation of ICB in PLWH (42) to boost immunity to reduce or eliminate the viral reservoir. However, clinical evidence on the impact of ICB as an HIV-1 cure intervention continues to be controversial (34,43–47). In this context, the simultaneous characterization of IRs co-expression and functional patterns across CD8+ T cells is critical to understanding Tex regulation in PLWH. This information is essential to identify novel targets for precise immunotherapies in PLWH on ART (48).

To address these questions, we performed supervised and unsupervised immunophenotypic analyses of IRs (PD-1, TIGIT, LAG-3, TIM-3, CD39), and functional markers (CD107a, IFNγ, and IL-2) across the landscape of CD8+ T cells over a decade of ART in PLWH and compared to PLWH with early infection and healthy individuals. We profile changes of bulk, SEB-activated, and HIV-1-specific CD8+ T-cells in PLHW and unfold a selective decrease of memory-like HIV-1-specific CD8+ clusters sharing TIGIT expression and low CD107a in PLWH on ART. Moreover, TIGIT blockade rescues CD107a expression without changes in IFNγ or IL-2 production on HIV-1-specific CD8+ T-cells. Of note, the response to TIGIT, TIM-3, or TIGIT+TIM-3 blockade was heterogeneous across HIV-1-specific CD8+ T-cell differentiation stages and functions, indicating the plasticity and complexity of the IRs pathways as targets for immune-base cure interventions.

## Results

### Alterations in CD8+ T-cell IRs frequencies and expression patterns in PLWH are not mitigated by ART

Although the co-expression of IRs is a hallmark of CD8+ Tex in HIV-1 infection (20–26), a detailed characterization of the combinatorial expression of IRs across CD8+ T-cell lineages in PLWH on long-term ART is still missing. To do this, we combined the analyses of IRs (PD-1, TIGIT, LAG-3, TIM-3, and CD39) and lineage markers (CD45RA, CCR7, and CD27) in CD8+ T cells from longitudinal samples in PLWH on ART by flow-cytometry (**Supplemental Figure 1**). We compare three groups; healthy controls (HC), PLWH with early infection (Ei), and PLWH on ART (S) with longitudinal samples available a median period of 2.2 (S1) and 10.1 (S2) years on fully suppressive ART (**Figure 1A, Source data Figure 1A**). The epidemiological and clinical characteristics of the study groups are detailed in **Supplemental Table 1** and **Source data Figure 1A**.

**Figure 1.**
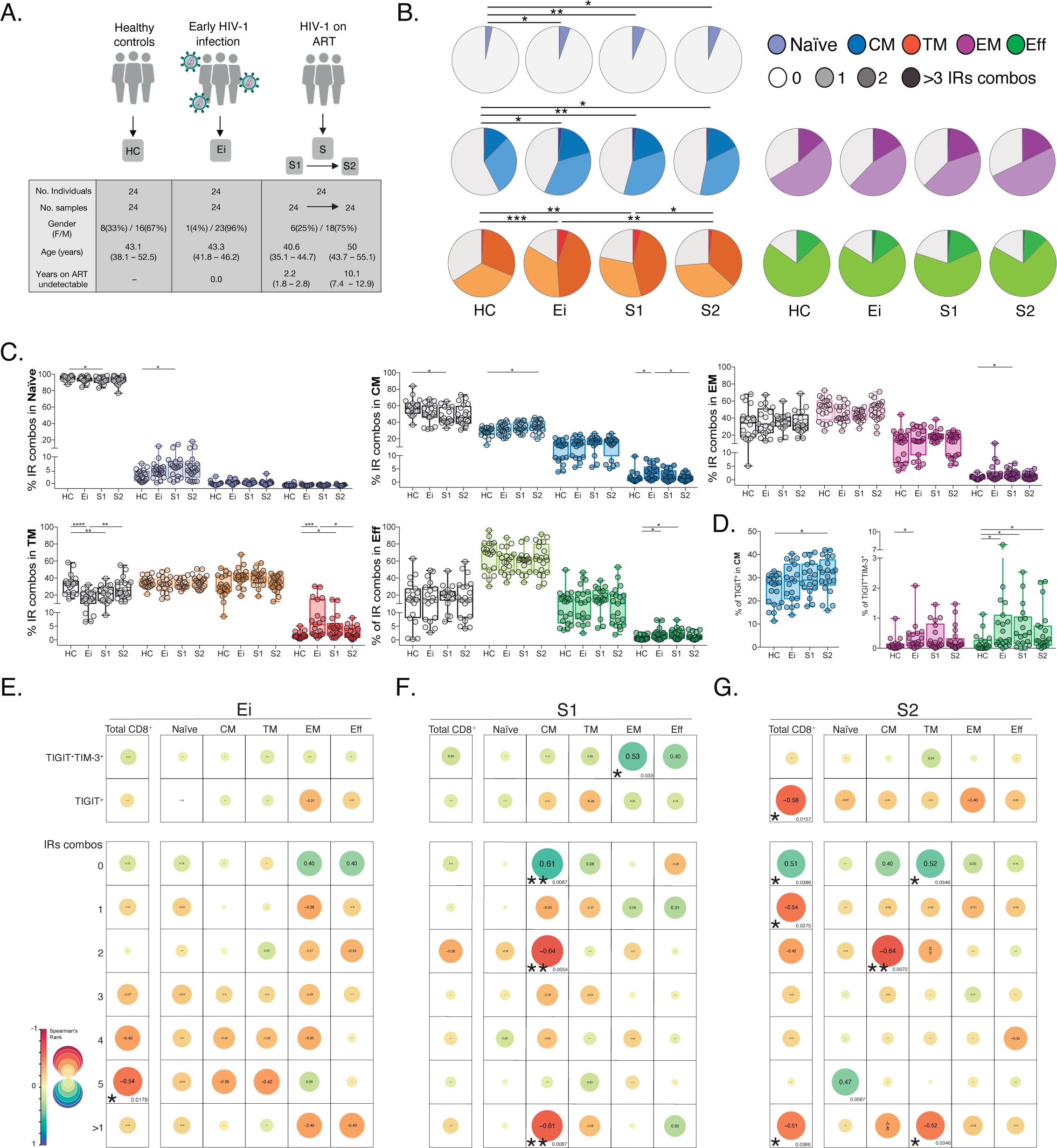
Patterns of IRs co-expression and correlations with CD4+ T -cell counts in PLWH. **(A)** Overview of study design and study groups, healthy controls (HC), PLWH in early HIV-1 infection (Ei), and PLWH on fully suppressive ART (S) in S1 and S2 time points. **(B)** The expression of IRs summarized in the pie chart is none, one, two, or more than three IRs expressed in CD8+ T-cell subsets. For statistical analysis, we used permutation tests using SPICE software. **(C)** Scatter plots showing the median and interquartile ranges of IR combinations in CD8+ T-cell subsets. **(D)** Scatter plots of the frequencies of single TIGIT^+^ expression in CM and TIGIT^+^TIM-3^+^ expression in EM and effector EFF CD8+ T cells. **(E-G)** Correlations between CD4+ T-cell counts as a function of TIGIT^+^, TIGIT^+^TIM-3^+^, and combinations of IRs from total CD8+ T cells and subsets in Ei (**E**), S1 (**F**), and S2 (**G**). The data in B to D represent the mean of two technical replicates. We used the Mann-Whitney U test for intergroup comparison (HC, Ei, S1, and S2) and the signed-rank test for intragroup comparison (S1 and S2). Holm’s method was used to adjust statistical tests for multiple comparisons. All possible correlations of the 32 Boolean IRs combinations are not shown. P-values: *<0.05, **<0.005 and ***<0.0005.

As shown in **Figure 1B (Source data Figure 1 B-C)**, we found persistent alterations in the expression and co-expression of IRs in naïve, central memory (CM), and transitional memory (TM) CD8+ T cells in PLWH on ART. These perturbations were maintained despite prolonged ART (S2) in naïve and CM, but not TM cells compared to HC. Deconvolution of IRs co-expression patterns by the number of receptors expressed (0, 1, 2, >3) further delineates a significant reduction of Naïve, CM, and TM CD8+ T cells lacking IRs expression concomitant with a significant increase of CD8+ T cells expressing one (Naïve and CM) or >3 IRs (TM) on ART (**Figure 1C**). Moreover, we observed an augment in effector memory (EM), TM, and effector (EFF) CD8+ T cells co-expressing >3 IRs in Ei and S1 that normalized in S2 (**Figure 1C**). Of note, out of the 32 possible combinations of IRs expression studied in CD8+ T-cell subsets (**Supplemental Figure 2C, Source data Figure S2**), single TIGIT^+^ expression in CM and dual TIGIT^+^TIM-3^+^ co-expression in EFF CD8+ T cells accounted for continuous increases in frequency under suppressive ART (**Figure 1D, Source data Figure 1D**). We confirmed similar increases in the frequency of TIGIT in total, CM, and TM CD8+ T cells on ART. These data contrast with transient changes in other IRs upon infection normalizing with ART (**Supplemental Figure 2A-B, Source data Figure S2**).

These initial findings led us to postulate associations between the expression of IRs in CD8+ T cells, persistent immune activation, and the degree of CD4+ T-cell immune recovery in PLWH on ART. For this purpose, we performed correlation analyses that revealed several negative associations between the frequency of CD8+ T cells expressing IRs and CD4+ T-cell counts across study groups (**Figure 1, E-G, Source data Figure 1E-G)**. Focusing on S1 (**Figure 1F**), we found significant negative correlations between CD4+ T-cell counts and frequencies of CM CD8+ T cells expressing 2 (p=0.0054, r=-0.64) and > 1 IRs (p=0.0087, r=-0.61). Focusing on S2 (**Figure 1G**), we observed significant negative correlations between CD4+ T-cell counts and the frequency of total CD8+ T cells expressing TIGIT^+^ (p=0.0157, r=-0.58), expressing 1 IRs (p=0.0386, r=-0.54) or >1 IRs (p=0.0386, r=-0.51). At the level of CD8+ T-cell subsets, the expression of 2 IRs in CM and >1 IR in TM negatively correlated with CD4+ T-cell counts (p=0.0072, r=-0.64; p=0.0346, r=-0.52, respectively) (**Figure 1G**). These correlations further indicate a negative relationship between IRs expression patterns and immune status in PLWH on long-term suppressive ART.

In summary, these data support changes in IRs expression not mitigated by long-term ART in total and CD8+ T-cell subsets expressing one or >1 IRs, particularly TIGIT^+^ and TIGIT^+^TIM-3^+^. These findings also uncover negative associations between IRs expression in CD8+ T cells and CD4+ T cell levels in PLWH on ART.

### Unsupervised phenotypic analyses of IRs across the CD8+ T-cell landscape in PLWH on ART

Next, to further characterize IRs expression across CD8+ T cells in PLWH on ART, we performed an unsupervised net-SNE analysis of flow-cytometry data. We concatenated 1,988,936 total CD8+ T cells and analyzed the phenotypes with the topographical regions of each surface marker tested (**Figure 2, A-B, Source data Figure 2A-D**). CD8+ T cells were classified into 38 cellular clusters distributed according to the relative marker expression of 14 parameters and represented using net-SNE and heatmaps (**Figure 2, C-D**).

**Figure 2.**
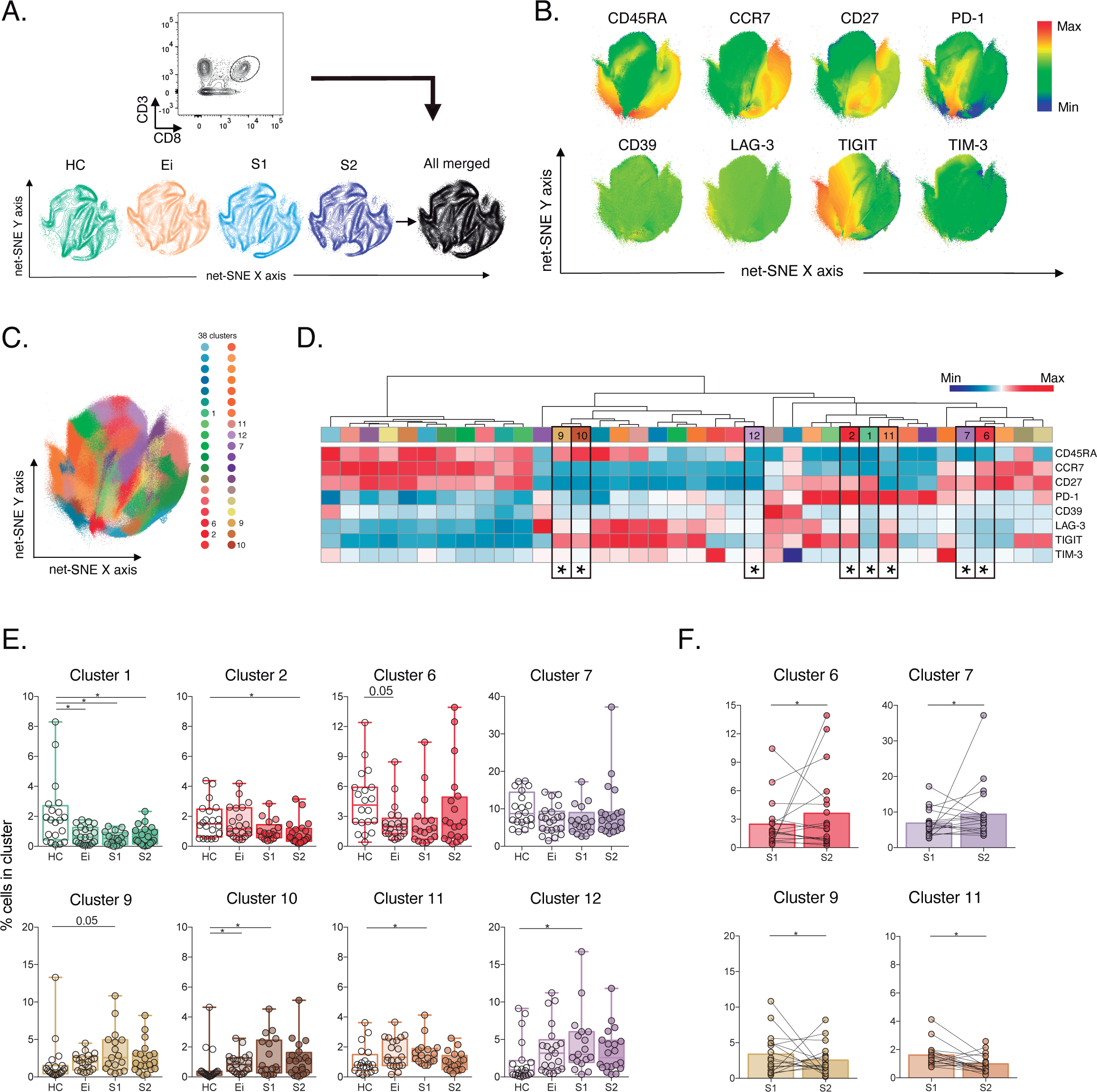
Unsupervised net-SNE analyses of total CD8+ T cells. **(A)** Gating strategy for selecting total CD8+ T cells (top), net-SNE plots of HC, Ei, S1, S2 and all merge groups. **(B)** Representative net-SNE visualization of surface markers. The colour gradient displays the relative marker expression. **(C)** Unsupervised KNN algorithm of 38 clusters coloured according to the legend. Only clusters with statistical differences are represented in the legend. **(D)** Heatmap of the median biexponential-transformed marker expression normalized to a -3 to 3 range of respective markers. Asterisks represent the clusters with statistical differences. **(E-F)** Scatter plots of intergroup (HC, Ei, S1 and S2) and intragroup (S1 and S2) cluster comparisons. Data represent the median and interquartile ranges of cluster cell frequency. We used the Mann-Whitney U test for intergroup analyses and the signed-rank test for intragroup analyses. Holm’s method was used to adjust statistical tests for multiple comparisons. P-values: *<0.05, **<0.005 and ***<0.0005.

Out of the 38 clusters identified, we found eight cellular clusters (#1, #2, #6, #7, #9, #10, #11, and #12) with significant differences by inter- and intragroup comparisons (**Figure 2D-E**). Most of the differentially expressed clusters shared memory-like phenotypes. Of note, clusters #6, #7, and #12 shared memory-like phenotypes and low expression of IRs. Meanwhile, clusters #9 and #10 shared effector-like phenotypes and co-expression of IRs, including TIGIT, LAG-3, and low TIM-3 (**Figure 2D**). Briefly, intergroup analyses demonstrated significant changes in composition and frequency with a decrease of #1, #6, and an increase of #10 in Ei compared with HC. Also, clusters #1 and #2 decreased in S1 and S2, respectively, and clusters #9, #10, #11, and #12 increased in S1 while tended to normalize in S2 compared with HC (**Figure 2E, Source data Figure 2 E-F**). Intragroup analyses further supported changes in cluster frequency and composition during long-term ART. Meanwhile, clusters #6 and #7 increased, and #9 and #11 decreased over time on ART (**Figure 2F**). These data support an expansion of memory-like clusters with low IR expression (#6 and #7) and a contraction of effector-like clusters sharing TIGIT expression (#9 and #11) during ART. Thus, unsupervised analyses support cellular clusters’ continuous expansion and contraction in frequency and composition across the landscape of CD8+ T cells in PLWH on ART.

### Unsupervised phenotypic characterisation of SEB-activated CD8+ T cells in PLWH on ART

Then, we evaluate CD8+ T-cell responses by bacterial superantigen activation with Staphylococcal enterotoxin B (SEB). Using SEB can provide complementary information on T-cell activation in response to pathogens involved in the disease by stimulating TCR-VB clonotypes (31,49,50). In this context, we analysed IRs expression and functional markers using unsupervised net-SNE analysis. We defined SEB-activated CD8+ T cells by the expression of at least one functional marker (CD107a, IFNγ, IL-2) upon incubation with SEB, as previously described (51–55) (**Figure 3, A-B, Source data Figure 3A-D**). We concatenated 253,021 SEB-activated CD8+ T cells and identified 29 unique clusters represented by net-SNE and heatmaps (**Figure 3, C-D**). Only six of the 29 clusters showed statistical differences by inter- and intragroup comparisons (**Figure 3D**). All differential clusters shared memory-like phenotypes (#2, #3, #5, #8, and #14) except cluster #6, with effector-like phenotype and low TIGIT expression. In addition, we observed a functional exclusion of clusters expressing IL-2 (#5 and #6) and those expressing CD107a and IFNγ (#2, #3, and #8) (**Figure 3D**). Intergroup comparisons identified increases in cluster #2 and a reduction of #14 and #5 with HIV-1 infection compared to HC. Additionally, ART was linked to the decrease of #5, #6, and #14 in S1 and #3 in S2 when compared with HC (**Figure 3E, Source data Figure 3 E-F**). Moreover, intragroup analyses identified an increase in clusters #6 and #14 and a reduction in #3 and #8 on ART. Of note, #6 characterizes by IL-2 expression in the without TIGIT expression. Meanwhile, clusters #3 and #8 express CD107a, IFNγ and variable expression of IRs (**Figure 3F**). In agreement with unsupervised clustering analyses, classical supervised analyses identified an augment of CD107a and IFNγ SEB-activated CD8+ T cells with Ei that normalized over time on ART. Also, we delineate significant increases of IL-2 SEB-activated CD8+ T cells across subsets and time on ART (**Supplemental Figure S3 A-C, Source data Figure S3**).

**Figure 3.**
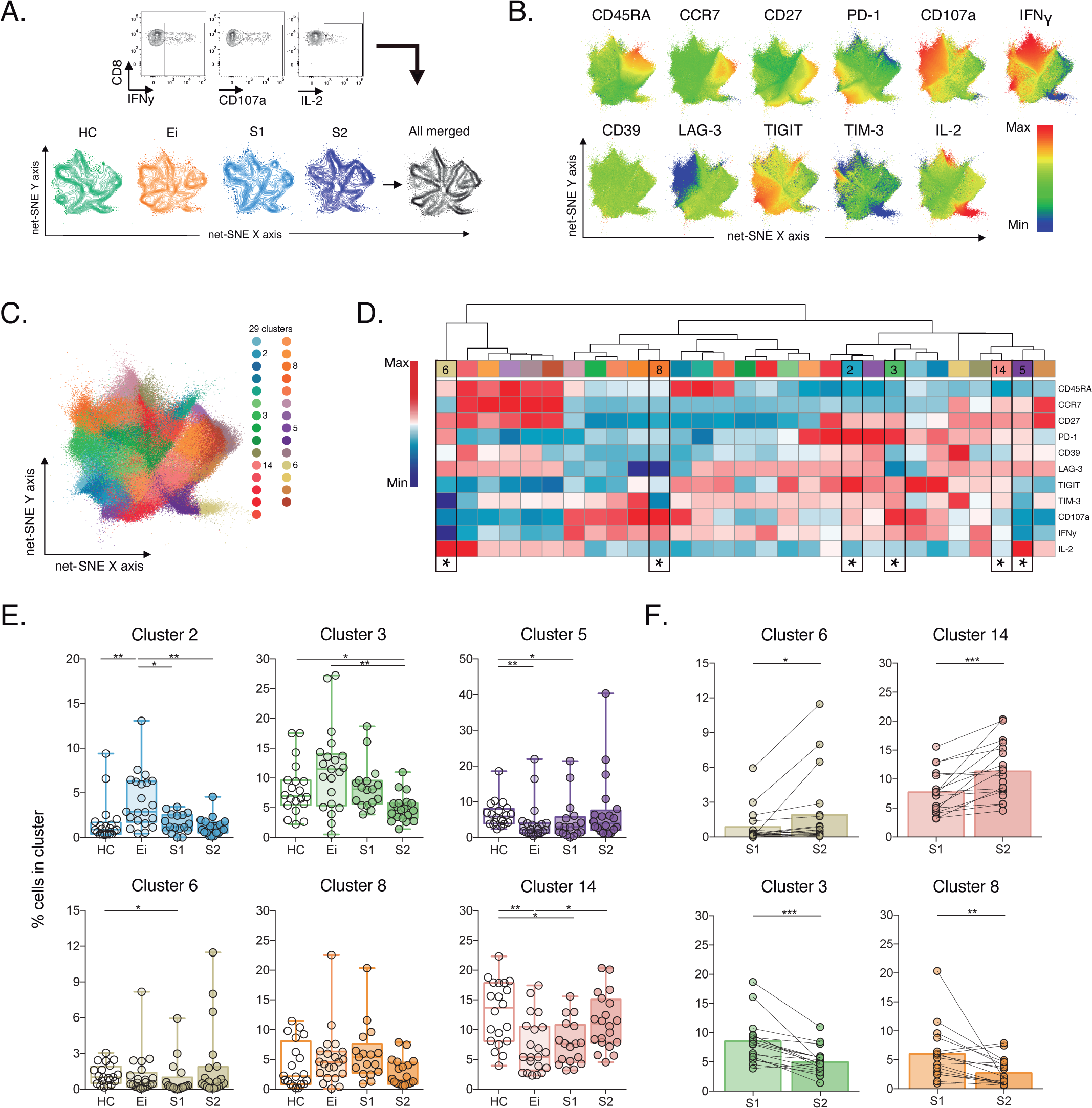
Unsupervised net-SNE analyses of SEB-activated CD8+ T cells. **(A)** Gating representation of CD107a, IFNγ and IL-2 expression in HIV-1-specific CD8+ T cells (top), net-SNE plots of HC, Ei, S1, S2 and merge groups of SEB-activated CD8+ T cells (bottom). **(B)** Representative net-SNE visualization of IR expression, lineage, and functional markers. The colour gradient displays relative marker expression. **(C)** Unsupervised KNN algorithm for 29 polyclonal clusters colour-coded according to the legend. Clusters with statistical differences between groups are represented in the legend. **(D)** Heatmap of the median biexponential-transformed marker expression normalized to a -3 to 3 range of respective markers. Asterisks represent the clusters with intergroup statistical differences. **(E-F)** Scatter plots of intergroup (HC, Ei, S1 and S2) and intragroup (S1 and S2) cluster comparisons. Data represent the median and interquartile ranges of cluster cell frequency. We used the Mann-Whitney U test for intergroup analyses and the signed-rank test for intragroup analyses. Holm’s method was used to adjust statistical tests for multiple comparisons. P-values: *<0.05, **<0.005, ***<0.0005.

These data support plasticity in the composition of SEB-activated CD8+ T-cell clusters with HIV-1 infection and ART. We observed the dominance of memory-like clusters with changes in composition and frequency in PLWH on ART, particularly IL-2 expression.

### Reduction of HIV-1-specific CD8+ T-cell clusters sharing memory-like phenotypes, TIGIT expression and low CD107a

Next, we characterize HIV-1-specific CD8+ T cells in PLWH on long-term ART compared to Early infected individuals (Ei), aiming to identify signatures of cellular dysfunction. We performed unsupervised net-SNE analyses in 53,751 cells concatenated, based on the production of at least one functional marker (CD107a, IFNγ, IL-2) in response to HIV-1-Gag peptides combined with lineage markers and IRs (**Figure 4, A-B, Source data Figure 4 A-D**). We defined 26 HIV-1-specific clusters by net-SNE analysis. Of note, only three showed significant differences between groups (#1, #2, and #3) (**Figure 4, C-D**). All three clusters decreased in frequency on ART and shared memory-like features (**Figure 4E, Source data Figure 4E**). Clusters #1 and #2 shared co-expression of IRs, mainly PD-1 and TIGIT, low CD107a, IFNγ, and higher expression of IL-2 than #3. Meanwhile, cluster #3 co-express (TIGIT, PD-1, and TIM-3) low expression of IL-2 and higher expression of CD107a and IFNγ (**Figure 4D**). Although CD107a, IFNγ, and IL-2 expression differed between clusters #1, #2, and #3, all shared TIGIT expression in the context of variable levels of TIM-3. These findings in memory-like clusters, together with the initial ones, accounting for increases in the frequency of TIGIT^+^ and TIGIT^+^TIM-3^+^ in CM and EFF cells on ART, led us to postulate then as potential markers of HIV-1-specific CD8+ T-cell dysfunction in PLWH on ART. Despite no significant changes observed in the total frequency of CD107a, IFNγ and IL-2 HIV-1-specific CD8+ T-cell responses between groups (**Supplemental Figure 4A-B**, **Source data Figure S4**). The analyses of TIGIT and TIGIT+TIM-3 HIV-1-specific memory subsets revealed decreases in the frequency of CD107a TIGIT HIV-1-specific CD8+ T cells limited to the CM compartment (**Figure 4F, Source data Figure 4F-G**). No changes in IFNγ and IL-2 were observed. Furthermore, polyfunctional analysis of TIGIT HIV-1-specific CM CD8+s identified a decrease in monofunctional CD107a^+^ as well as in bifunctional CD107a^+^IFNy^+^ and CD107^+^IL-2^+^ cells overtime on ART (**Figure 4G**). Overall, these data support a reduction of HIV-1-specific CD8+ T-cell clusters sharing memory-like phenotypes, TIGIT expression and low CD107a in PLWH on ART.

**Figure 4.**
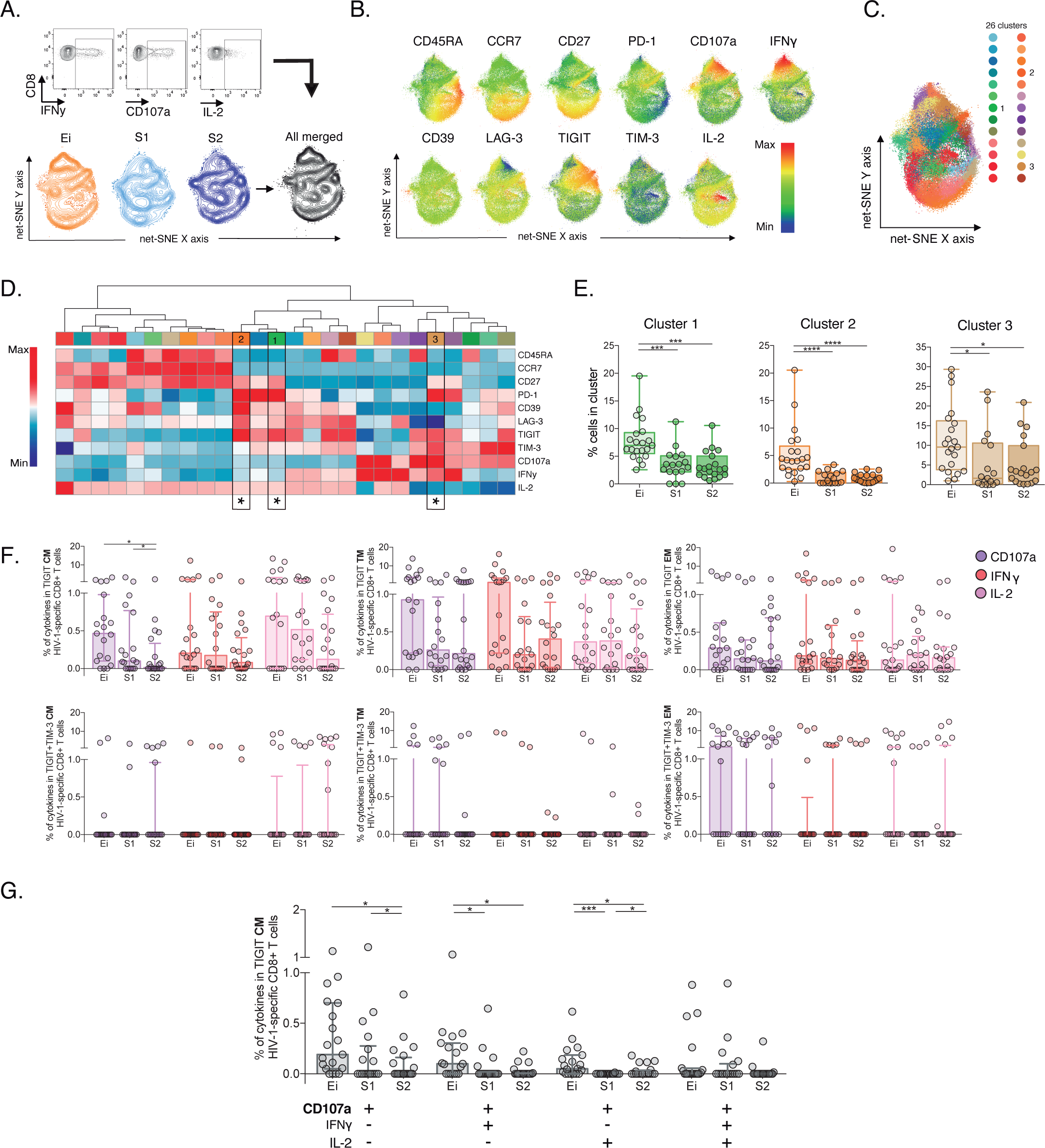
Unsupervised and supervised analyses of HIV-1-specific CD8+ T cells. **(A)** Gating representation of CD107a, IFNγ and IL-2 expression in HIV-1-specific CD8+ T cells (top), net-SNE plots of Ei, S1, S2 and merge groups for HIV-1-specific CD8+ T cells (bottom). **(B)** Representative net-SNE plots for surface and functional markers. The colour gradient displays relative marker expression. **(C)** Unsupervised KNN algorithm for 26 HIV-1-specific clusters colour-coded according to the legend. Only clusters with statistical differences are represented in the legend. **(D)** Heatmap of the median biexponential-transformed marker expression normalized to a -3 to 3 range of respective markers. Asterisks represent the clusters with intergroup statistical differences. **(E)** Scatter plots of intergroup (Ei, S1 and S2) cluster comparisons with significant statistical differences. Data represent the median and interquartile ranges of cluster cell frequency. **(F)** CD107a, IFNγ, and IL-2 frequency of expression in TIGIT+ (upper panel) and TIGIT+TIM-3+ (bottom panel) HIV-1-specific memory CD8+ T cell subsets. Scatter plots represent the median and interquartile ranges. **(G)** Polyfunctional analyses of CD107a IFNγ and IL-2 expression in CM TIGIT HIV-1-specific CD8+ T cells. Scatter plots represent median and interquartile ranges. We used the Mann-Whitney U test for intergroup analyses and the signed-rank test for intragroup analyses. Holm’s method was used to adjust statistical tests for multiple comparisons. P-values: *<0.05, ***<0.0005, and ****<0.0001.

### TIGIT blockade restores CD107a expression but not IFNγ or IL-2 production in HIV-1-specific CD8+ T cells

Then, we decided to explore the blockade of TIGIT and TIM-3 pathways as targets for the recovery of CD107a and potentially IFNγ and IL-2 production in HIV-1-specific CD8+ T-cells. We performed short-term ICB experiments using monoclonal antibodies αTIGIT, αTIM-3, and αTIGIT+ αTIM-3 in PBMC from PLWH on suppressive ART (S) with previous immunophenotype.

After short-term ICB, we monitored changes in CD107, IFNγ, and IL-2 in total and subsets of HIV-1-specific CD8+ T cells by flow cytometry (**Figure 5A)**. The Net-SNE TIGIT and TIM-3 projections are represented in **Figure 5B (Source data Figure 5B)**. At the level of total HIV-1-specific CD8+ T-cells, short-term ICB experiments demonstrate a specific increase of CD107a expression by αTIGIT (isotype vs. αTIGIT; p<0.05) and αTIGIT + αTIM-3 (isotype vs. αTIGIT + αTIM-3; p<0.05) blockade (**Figure 5B, left**). Moreover, the recovery of CD107a by αTIGIT was consistent across HIV-1-specific CD8+ T-cell subsets, being particularly marked for CM (isotype vs. αTIGIT; in CM p<0.005) (**Figure 5C**) in agreement with our previous findings. Of note, αTIM-3 blockade did not show any effect, and dual blockade of αTIM-3+αTIGIT did not reveal an additive effect (**Figure 5C**). No changes in IFNγ and IL-2 production were observed for any conditions tested (**Figure 5, D-E, Source data Figure 5 D-E**).

**Figure 5.**
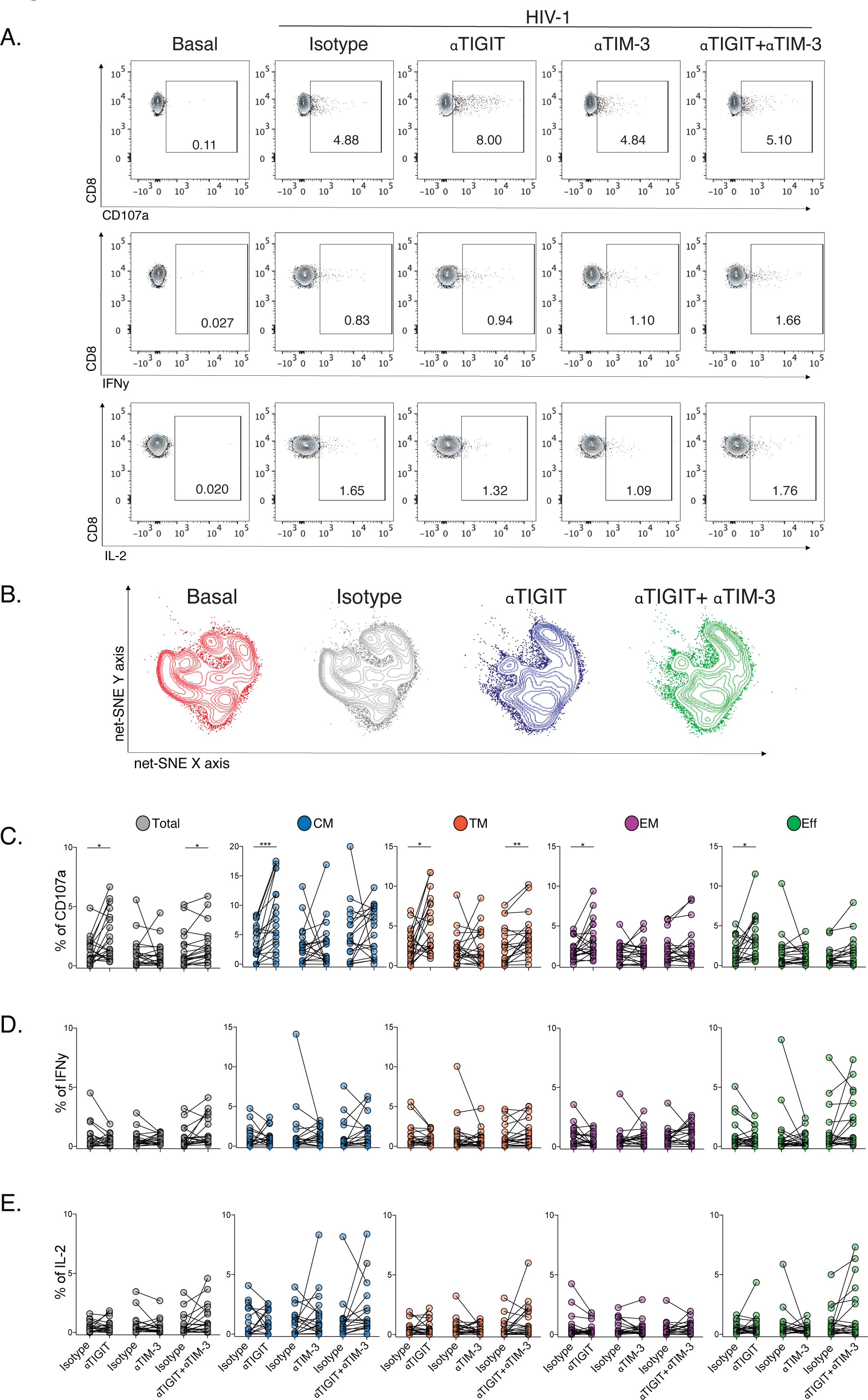
Effect of TIGIT, TIM-3 and TIGIT+TIM-3 mAb blockade in HIV-1-specific CD8+ T cell responses in PLWH on ART. **(A)** Representative flow cytometry plots gated on CD8+ T-cells, in the absence of HIV-1 Gag stimulation (basal condition) and presence of HIV-1 Gag stimulation with isotype control, αTIGIT, αTIM-3, and αTIGIT+αTIM-3 antibodies for CD107a, IFNγ and IL-2 expression. **(B)** Representative net-SNE plots for HIV-1-specific CD8+ T cells from PLWH concatenated and merged according to the condition. **(C-E)** Frequency of CD107a, IFNγ, and IL-2 expression in total and HIV-1-specific CD8+ T cell subsets for the various conditions tested. The Wilcoxon matched-pairs signed ranked test calculated statistical differences. The data represent the mean of two technical replicates. P-values: =0.05, *<0.05, **<0.005 and ***<0.0005.

These data support heterogeneity in the functional recovery of HIV-1-specific CD8+ T-cells by differentiation stage based on αTIGIT, αTIM-3, and αTIGIT+αTIM-3 blockade. Overall, these results identify the targeting of TIGIT to recover the degranulation in HIV-1-specific CD8+ T cells, particularly within the CM compartment in PLWH on ART.

## Discussion

CD8+ Tex displays a range of functional defects in PLWH early in HIV-1 infection and during ART (16–19). The expression of IRs is a hallmark of Tex (20,22,23) (14), and co-expression of IRs has been associated with HIV-1 disease progression (15,25,29,30,32) and cancer severity (56–58). Although ICB has demonstrated promising results in cancer remission (40,41), its applicability in HIV-1 as a cure intervention remains unclear (34,43–47). Therefore, it is essential to understand the patterns of IRs expression and function across the CD8+ T-cell landscape to identify targets for ICB broadly applicable to PLWH on ART (48).

Here, we immunophenotype CD8+ T cells using five IRs and three functional markers in PLWH over the ten years of ART. In this way, we overcome previous study limitations based on single IRs expression, bulk CD8+ T cells, and cross-sectional data (20,23,25,29,32). Our results demonstrated a marked and significant increase in TIGIT CD8+ T cells, particularly within the central memory compartment, not ameliorated by long-term ART. Also, TIGIT CD8+ T cells negatively correlated with CD4+ counts in PLWH on ART. These data support continuous expression of TIGIT despite ART in agreement with previous studies (24,25,30,57,59) and uncover novel associations between TIGIT expression in CD8+ T cells and poorer immune status in PLWH on ART. Thus, these data indicate a specific contribution of TIGIT expression to persistent immune activation and poor CD4+ recovery on ART.

In contrast, we observed transient increases of PD-1, LAG3, TIM3 and CD39 expression in total, memory and effector CD8+ T cells that normalize over time on ART. The potential biological implications of such a difference may relate to the nature of each receptor and the specific immune regulatory pathway activated during HIV-1 infection and ART. Also, these divergences may be influenced by the presence of γ-chain cytokines, such us IL-2, IL-15 and IL-21, in plasma able to upregulate the TIGIT expression in CD8+ T cells (25). A previous study demonstrated an association between high levels of IL-15 and high TIGIT expression in CD4+ T cells with suboptimal CD4+ recovery in PLWH on ART (60).

Our study combined high-dimensional supervised and unsupervised analysis providing an unprecedented deep immunophenotype of the CD8+ T-cell landscape in PLWH over a decade on ART. We explored complementary levels of complexity for the characterization of CD8+ T cells, including an absence of stimuli (bulk), the presence of antigen-independent stimuli (SEB), and the presence of antigen-specific stimuli (HIV-1).

In the absence of stimuli, supervised analyses confirmed heterogeneous and complex patterns of IRs co-expression across CD8+ T cell lineages altered by HIV-1 infection and shaped by ART (16,21,30,61,62). Furthermore, unsupervised analyses added complexity to previous data by delineating in profound detail early and continuous changes of contraction and expansion of cellular clusters with HIV-1 infection and time on ART. We tracked memory-like expansion and effector-like cluster contraction over time on ART according to the establishment of memory responses. Similarly, in the presence of antigen-independent stimuli, we identified continuous changes in cellular cluster composition by contraction and expansion of CD8+ cellular cluster with infection and treatment. We observed a dominance of memory-like clusters with changes in composition and frequency tracked by an augment of IL-2 expressing clusters on ART both by supervised and unsupervised analyses. The IL-2 expression regulates proliferation and homeostasis and contributes to the generation of long-term memory responses (31,49,50), suggesting a partial functional remodelling of CD8+ T-cell populations independent of antigen in PLWH over time on ART. Thus, our findings support the contribution of IRs co-expression in CD8+ Tex and T-cell activation favouring a continuous reshaping of memory and effector-like CD8+ cellular clusters. These findings indicate the enormous plasticity and constant homeostasis of CD8+ T cells in PLWH during a decade of ART (63).

In the presence of HIV-1-specific stimuli, supervised and unsupervised analyses delineate a reduction of HIV-1-specific CD8+ T cells sharing memory phenotypes, TIGIT expression and low CD107a. Our findings focused on the memory compartment of TIGIT expressing HIV-1-specific CD8+ T cells demonstrating a decrease in monofunctional CD107a^+^ and bifunctional CD107a^+^IFNγ^+^ and CD107a^+^IL-2^+^ cells. Previous studies support a direct correlation between monofunctional CD107a^+^ and Eomes intensity in exhausted HIV-1-specific CD8+ T cells (17). Although this study did not include the expression of TIGIT across analyses, it may suggest an association between TIGIT expression and the T-bet /Eomes axis in charge of regulating the exhaustion and memory cell fate of CD8+ T cells (64).

Our findings further support the role of TIGIT as a signature of dysfunctional and Tex antiviral responses (25). Indeed, the blockade of the TIGIT pathway restored CD107a expression in HIV-1-specific CD8+ T cells across cellular compartments with a marked effect in the central memory compartment according to our findings from HIV-1-specific immunophenotype of TIGIT CD8+ T cells. To our knowledge, our study is the first to demonstrate the recovery of CD107a expression in HIV-1-specific CD8+ T cells by TIGIT blockade. These data contrast with Chew et al., which observed the recovery of IFNγ production by TIGIT blockade. However, they also reported a reduction in CD107a expression in TIGIT+ CD8 T cells in response to aCD3/aCD28 activation, supporting a dysfunctional profile of TIGIT+ CD8+ T cells. Differences between study groups accounting for time on ART, samples tested and interindividual variability of in vitro ICB experiments may account for some of the differences observed.

Although the mechanistic behind TIGIT signalling and CD107a expression are not fully understood, low CD107a expression has been linked to the terminal T-bet^dim^Eomes^hi^ exhausted phenotype, and HIV-1-specific CD8+ T cells expressing TIGIT can degranulate to a certain extent (17,24). Moreover, our data did not support an additive effect recovering HIV-1-specific CD8+ T cells function of TIGIT and TIM-3 combinatorial blockade over TIGIT blockade. The redundancy and promiscuity of TIM-3 for several ligands, including Gal-9, CEACAM-1, PtdSer, and HMGB-1 (65,66), may be associated with these results. We cannot exclude the impact of TIGIT and TIM-3 blockade in other cell types (57,58,67).

We acknowledge several study limitations; First, the sample size of study groups and the use of peripheral blood samples underestimate the potential contribution of TIGIT expression to Tex in lymphoid tissues. Second, the use of only Gag as stimuli for the characterization of HV-1-specific CD8+ T cell responses in the absence of TCR sequencing. Using alternative HIV-1 antigens such as Nef, Env or Pol may provide additional information on the profile of CD8+ T-cell functional responses against early and late-expressed viral proteins in PLWH on ART (68, 69). Third, limited ICB experiments to CD107a, INFγ and IL-2 functional markers without complementary cytotoxic markers (perforin, granzyme B). Forth, complementary transcriptomic, epigenetic, and metabolic markers are needed for a complete description of Tex’s immune signatures linked to TIGIT expression in HIV-1 specific CD8+ T cells in PLWH on ART.

In summary, our study profile with unprecedented detail continuous reshaping of memory-like and effector-like of CD8+ T cellular clusters in PLWH over a decade of ART. The study identifies the TIGIT as a critical target for T-cell dysfunction, and Tex associated the loss of CD107a expression in HIV-1-specific CD8+ T cells. These findings support targeting the TIGIT/CD155 axis for Tex precision immune-base curative interventions in PLWH at all ART stages.

## Materials and Methods

### Study groups

This retrospective study analyzed clinical data and biological sample availability from 3,000 patients assigned to the HIV-1 clinical unit of the Germans Trias i Pujol University Hospital. We included individuals with cryopreserved PBMCs available in our collection. We identified 24 chronically HIV-1-infected individuals who had been treated mainly with a combination of NNRTI and NRTI for more than ten years with sustained virological suppression (<50 HIV-1-RNA copies/ml) and with longitudinal biological samples at timepoint 1 (S1), 2.2 (1.8 - 2.8) years undetectable on ART, and at time point 2 (S2), 10.1 (7.4 – 12.9) years undetectable on ART (**Supplemental Table 1, Figure 1A, Source data Figure 1A**). We excluded individuals with integrase inhibitors, ART as monotherapy, and treatments with mitochondrial toxicity, including Trizivir, d4T, ddI, AZT and blips over the ART period (S1-S2) to ensure homogeneous treatment over time. For comparative purposes, we included 24 early HIV-1-infected individuals (Ei) defined in a window of 1.3 (0.77 - 17.8) weeks after seroconversion in the absence of ART and 24 healthy controls (HC). The groups were balanced by age to the S2 samples to avoid confounding effects on IR expression.

### CD8+ T cell immunophenotype

Cryopreserved PBMCs from the study groups were thawed and rested overnight at 37°C in a 5% CO_2_ incubator. The following day, PBMCs were incubated for six hours at 37°C in a 5% CO_2_ incubator under RPMI complemented medium 10% FBS in the presence of CD28/49d co-stimulatory molecules (1μl/ml, BD), Monensin A (1μl/ml, BD Golgi STOP), and anti-human antibody for CD107a (PE-Cy5, clone H4A3, Thermo Fisher Scientific). PBMCs were left unstimulated, stimulated with SEB (1μg/ml, Sigma-Aldrich), and stimulated with HIV-1-Gag peptide pool (2μg/peptide/ml, EzBiolab). After six hours of stimulation, cells were rested overnight at 4°C as previously described (34). The next day, PBMCs were washed with PBS 1X and stained for 25 minutes with the Live/Dead probe (APC-Cy7, Thermo Fisher Scientific) at RT to discriminate dead cells. Cells were washed with PBS 1X and surface stained with antibodies for 25 minutes at RT. We used CD3 (A700, clone UCHT1, BD), CD4 (APC-Cy7, clone SK3, BD), CD8 (V500, clone RPA-T8, BD), CD45RA (BV786, clone HI100, BD), CCR7 (PE-CF594, clone 150503, BD), CD27 (BV605, clone L128, BD), TIGIT (PE-Cy7, clone MBSA43, Labclinics SA), PD-1 (BV421, clone EH12.1, BD), LAG-3 (PE, clone T47-530, BD), TIM-3 (A647, clone 7D3, BD) and CD39 (FITC, clone TU66, BD) antibodies. Afterwards, cells were washed twice in PBS 1X, fixed, and permeabilized with Fix/Perm kit (A and B solutions, Thermo Fisher Scientific) for intracellular cytokine staining with anti-human antibodies of IFNγ (BV711, clone B27, BD) and IL-2 (BV650, clone MQ1-17H12, BD). Finally, stained cells were washed twice with PBS 1X and fixed in formaldehyde 1%.

### TIGIT, TIM-3 and TIGIT+TIM-3 short-term checkpoint blockade

We selected cryopreserved PBMCs from S1 (n=10) and S2 (n=10). Samples were previously characterized by the expression of TIGIT and TIM-3 on total CD8+ T cells. PBMCs were thawed and rested for four hours at 37°C in a 5% CO_2_ incubator. Next, cells were incubated under RPMI complemented medium 10% FBS with 1μl/ml of anti-CD28/CD49d and 1μl/ml of Monensin A overnight at 37°C in a 5% CO_2_. PBMCs are divided in the following conditions; 1) unstimulated, 2) SEB (1μg/ml, Sigma-Aldrich) and 3) HIV-1-Gag peptide pool (2μg/peptide/ml) in the absence or presence of αTIGIT and/or αTIM-3, and its respective isotype antibodies. For the single blockade of TIGIT (αTIGIT), we included Ultra-LEAF™ purified anti-human TIGIT antibody (10μg/ml, clone A15153G, Biolegend) or its control isotype Ultra-LEAF™ purified mouse IgG2a antibody (10μg/ml, MOPC-173, Biolegend). For single TIM-3 blockade (αTIM-3), we used Ultra-LEAF™ purified anti-human TIM-3 antibody (10μg/ml, clone F38-2E2, Biolegend) or its respective isotype Ultra-LEAF™ purified mouse IgG1 antibody (10μg/ml, MOPC-21, Biolegend). Finally, we included αTIGIT+αTIM-3 or their respective IgG2+IgG1 isotypes for a combinational blockade. The next day, PBMCs were surface and intracellularly stained with the panel of antibodies and the methodology described in the section above.

### Supervised immunophenotype data analysis

Stained PBMCs were acquired on an LSR Fortessa cytometer using FACSDiVa software (BD). Approximately 1,000,000 events of PBMCs were recorded per specimen. Antibody capture beads (BD) were used for single-stain compensation controls. Flow cytometry data were analyzed with FlowJo software v10.6.1, and fluorescence minus one (FMO) was used to set manual gates. We analyzed CD8+ T cells by excluding dump and CD4+ T cells. We excluded patients with <20% viability in lymphocytes and total CD8+ T cells (**Source data Figure 1A**). As previously described, we measured by supervised and classical analyses the IRs expression in CD8+ T-cell subsets, including Naïve, central memory, transitional memory, effector memory and effector CD8+ T cells (20,34). We performed two technical replicates for SEB-activated and HIV-1-specific CD8+ T-cell cytokine production. We considered the cytokine response positive after background subtraction (mean of two technical replicates) used as the cut-off value. For each independent sample, we recorded a median of 1,000 events and 50 events positive for cytokines for total and CD8+ T cell subsets, respectively.

### Unsupervised immunophenotype data analysis

The phenotypic and functional characterization of cellular populations was analyzed by using t-Distributed Stochastic Neighbor Embedding (t-SNE) (70) and net-SNE (71) dimensionality reduction algorithms to visualize single-cell distributions in two-dimensional maps. Briefly, cell intensity was z-normalized, and a randomly selected subset of cells, at least 1,000 cells per sample, was passed through the t-SNE algorithm. The resulting t-SNE dimension was then used to predict the position of all remaining CD8+ T cells acquired per sample from each group using the net-SNE algorithm based on neural networks. For functional analysis, we selected polyclonally activated and HIV-1-specific CD8+ T cells producing at least one cytokine CD107a, IFNγ, or IL-2 under SEB or HIV-1 conditions, respectively. In parallel, we discovered cell communities using the Phenograph clustering technique. It operates by computing the Jaccard coefficient between nearest neighbours, which was set to 30 in all executions, and then locating cell communities (or clusters) using the Louvain method. The method creates a network indicating phenotypic similarities between cells. The netSNE maps included representations of the identified cell communities, and additionally, we built a heatmap with the clusters in the columns and the markers of interest in the rows to better comprehend the phenotypical interpretation of each cluster. The colour scale displays each marker’s median intensity on a biexponential scale. We calculated quantitative assessments of cellular clusters in the percentage of cells for each sample to analyze and compare the distribution between HC, Ei, S1, and S2 groups, similar to the classical flow cytometry analysis.

### Statistics

Bivariate analysis was conducted using nonparametric methods as follows: Mann-Whitney U test for independent median comparison between groups, Wilcoxon signed-rank test for paired median changes over time, permutation test for composition distribution between groups, Kruskal-Wallis test for comparison between more than two groups, and spearman linear correlation coefficient to study the association between continuous variables. Holm’s method was used when appropriate to adjust statistical tests for multiple comparisons with a significance level of 0.05. All statistics and single-cell analyses were conducted using the R statistical package (72). The selection of clusters was performed through the significant differences obtained by inter- and intragroup comparisons between groups. Moreover, pattern distribution and graphical representations of all possible Boolean combinations for IRs co-expression and functional markers were conducted using the data analysis program Pestle v2.0 and SPICE v6.0 software (73). Graph plotting was performed by GraphPad Prism v8.0 software and R packages. The data represents the mean of two technical replicates.

### Study Approval

The study was conducted according to the principles of the Declaration of Helsinki (Fortaleza, 2013). The Hospital Germans Trias i Pujol Ethics Committee approved all experimental protocols (PI14-084). For the study, subjects provided written informed consent for research purposes of biological samples taken from them.

## Author contributions

OB-L and JGP conceptualized and designed the experiments. OB-L, EJ-M, and RP developed immunophenotypic analyses. OB-L, MAM, and JC developed short-term antibody blockade experiments. OB-L, DO, MAM, AP, AT, AS, R-PS, and JGP performed supervised and unsupervised analyses. JD, JRS, BC, and JGP recruited the study participants. OB-L and JGP wrote the manuscript. All authors revised it critically for important intellectual content and have approved the final version submitted for publication.

## Acknowledgements

We thank the Flow Cytometry Core Facility from the Germans Trias i Pujol Research Institute (IGTP). This research was supported by National Health Institute Carlos III grant PI17/00164. Oscar Blanch-Lombarte was funded by an AGAUR-FI_B 00582 PhD fellowship from Catalan Government and the European Social Fund. Redes Temáticas de Investigación en SIDA (ISCIII RETIC RD16/0025/0041) funded Esther Jiménez-Moyano. This study has received partial funding from Grifols. The funders had no role in study design, data collection, analysis, or the decision to publish or draft the manuscript.

## Supplemental Figures

**Supplemental Figure 1.**
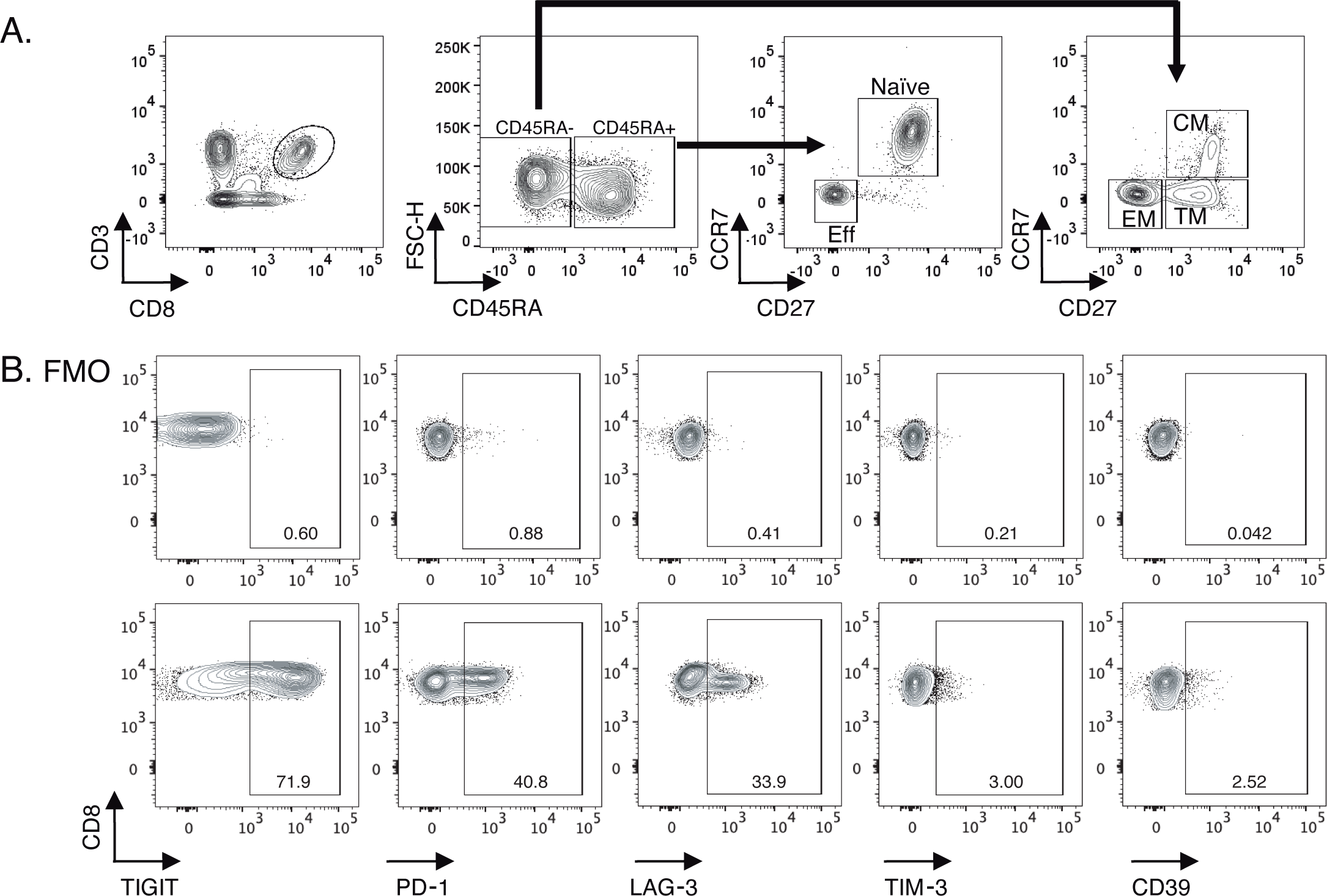
Gating strategy for identifying total CD8+ T-cell subsets and IRs expression levels. **(A)** Zebra dot plots showing the gating strategy used for the identification of total CD8+ and cellular using lineage markers (CD45RA, CCR7, and CD27) subsets as follow; naïve (CD45RA+ CCR7+ CD27+), Eff (CD45RA+ CCR7-CD27-), CM (CD45RA-CCR7+ CD27+), TM (CD45RA-CCR7-CD27+) and EM (CD45RA-, CCR7-, and CD27-). **(B)** Zebra dot plots of FMO and IRs staining demonstrating TIGIT, PD-1, LAG-3, TIM-3, and CD39 expression in cryopreserved PBMCs.

**Supplemental Figure 2.**
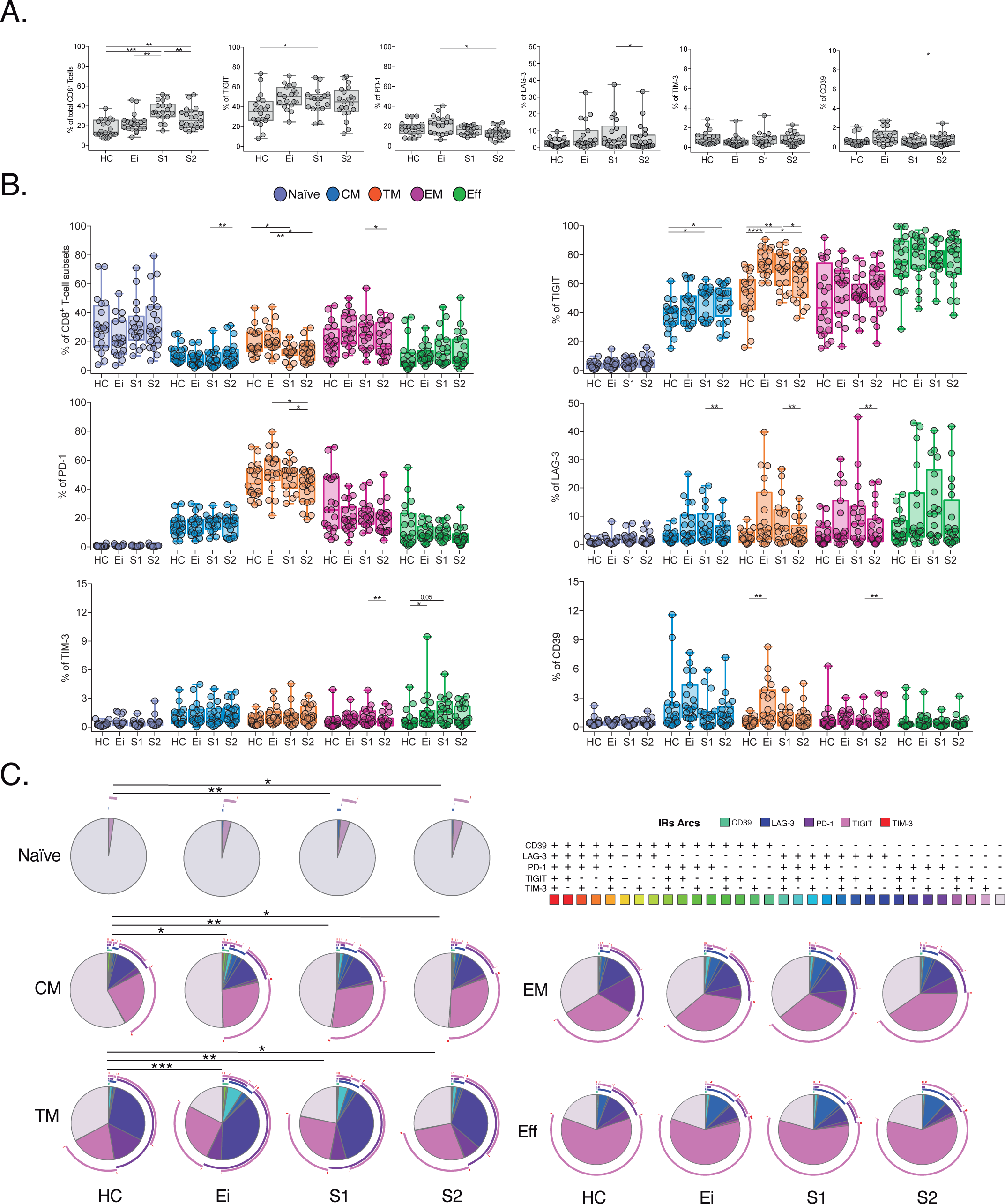
Expression profile of IRs in total and CD8+ T cell subsets across study groups. **(A)** Frequency of CD8+ T cells and IRs expression in total CD8+ T cells across study groups (HC, Ei, S1 and S2) **(B)** Frequency of IRs expression in CD8+ T-cell subsets. Scatter plots represent median and interquartile ranges. **(C)** Co-expression profile of IRs across CD8+ T cell subset. Pie charts represent the frequency for the 32 possible combinations for TIGIT, PD-1, LAG-3, TIM-3, and CD39 expression. The pie charts represent the frequency of expression according to the colour code, and the arcs indicate the frequency of each IR. We used permutation tests of SPICE software for statistical analysis of IRs co-expression data. We used the Mann-Whitney U test for intergroup analyses and the signed-rank test for intragroup analyses. Holm’s method was used to adjust statistical tests for multiple comparisons. P-values: *<0.05, **<0.005, ***<0.0005, ****<0.00001.

**Supplemental Figure 3.**
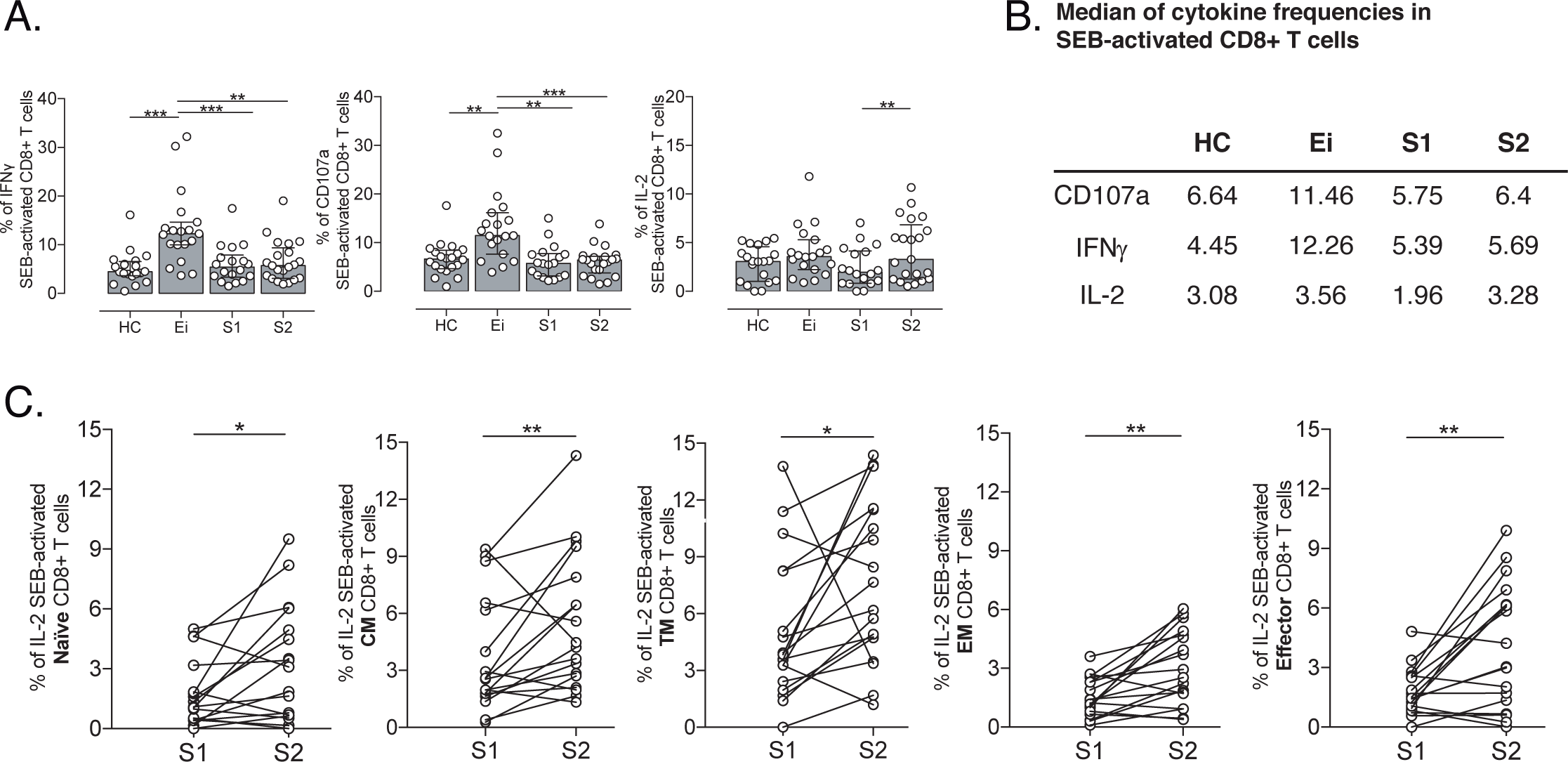
Supervised analyses of SEB-activated CD8+ T cells. (**A**) The total frequency of CD107a, IFNγ and IL-2 SEB-activated CD8+ T cells in HC, Ei, S1 and S2. Data represent median and interquartile ranges. (**B**) Table of median frequencies of CD107a, IFNγ and IL-2 SEB-activated CD8+ T-cell. (**C**) Frequency of IL-2 SEB-activated CD8+ T cell subsets in S1 and S2. We used the Mann-Whitney U test for independent median intergroup comparison and the signed-rank test for paired median changes between S1 and S2. The Holm’s method was used to adjust statistical tests for multiple comparisons. P-values: *<0.05, **<0.005, ***<0.0005.

**Supplemental Figure 4.**
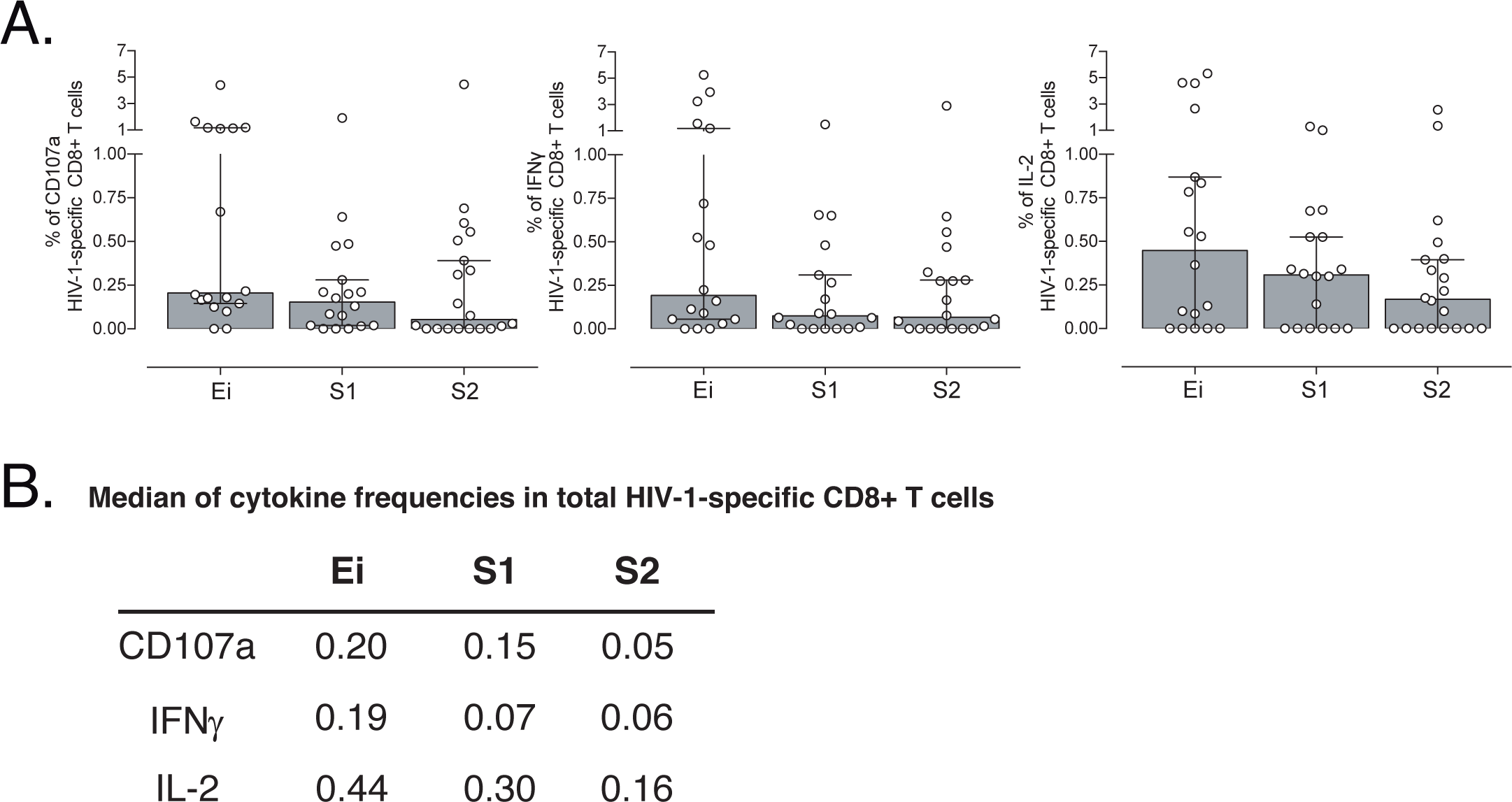
Supervised analyses of HIV-1-specific CD8+ T cell responses **(A)** Frequency of CD107a, IFNγ and IL-2 HIV-1-specific CD8+ T cells in HC, Ei, S1 and S2. Data represent median and interquartile ranges. (**B**) Table of median frequencies of CD107a, IFNγ and IL-2 HIV-1-specific CD8+ T cells. We used the Mann-Whitney U test for intergroup analyses and the signed-rank test for intragroup analyses. Holm’s method was used to adjust statistical tests for multiple comparisons.

**Supplemental Table 1.** Epidemiological characteristics of the study groups.

|  | Healthy Control | Early HIV-1 infection | HIV-1 on ART |  | p |
| --- | --- | --- | --- | --- | --- |
| Study Groups | HC | Ei | S |  |  |
| Number of subjects | 24 | 24 | 24 |  |  |
| Gender (M/F)<br>[n (%)] | 16/8<br>(67/33%) | 23/1<br>(96/4%) | 18/6<br>(75/25%) |  | 0.091 |
| Weeks since seroconversion<br>[median (IQR)] | - | 1.3<br>(0.77 - 17.8) | - | - |  |
| Number of samples | 24 | 24 | 24 (S1) | 24 (S2) |  |
| Age at sample (years)<br>[median (IQR)] | 43.1<br>(38.1 - 52.5) | 43.3<br>(41.8 - 46.2) | 40.6<br>(35.1 - 44.7) | 50.0<br>(43.7 - 55.1) | 0.0035 |
| CD4+ T-cell counts at sample per mm<sup>3</sup><br>[median (IQR)] | - | 630<br>(557 - 757.3) | 573<br>(464 - 992) | 910<br>(799.8 - 1111) | 0.0001 |
| Years undetectable at sample on ART<br>[median (IQR)] | - | 0.0 | 2.2<br>(1.8 - 2.8) | 10.1<br>(7.4 - 12.9) | <0.0001 |
| VL log copies/ml at sample [median (IQR)] | - | 4.5 (4.3 - 5.1) | undetectable | undetectable | <0.0001 |

Abbreviations: IQR, interquartile range; P-values were calculated using Kruskal-Wallis test (Dunn's correction) or X<sup>2</sup>-test.

## Notes

### Competing Interest Statement

The authors have declared no competing interest.

### Summary of Updates

Based on comments for reviews

